# Vacuolar sucrose homeostasis is critical for development, seed properties and survival of dark phases of Arabidopsis

**DOI:** 10.1101/2020.01.20.912253

**Authors:** Duc Phuong Vu, Cristina Martins Rodrigues, Benjamin Jung, Garvin Meissner, Patrick A.W. Klemens, Daniela Holtgräwe, Lisa Fürtauer, Thomas Nägele, Petra Nieberl, Benjamin Pommerrenig, H. Ekkehard Neuhaus

## Abstract

Although we know that most of the cellular sucrose is present in the cytosol and vacuole, our knowledge on the impact of this sucrose compartmentation on plant properties is still fragmentary. Here we attempted to alter the intracellular sucrose compartmentation of Arabidopsis mesophyll cells by either, overexpression of the vacuolar sucrose loader *Bv*TST2.1 or by generation of mutants with decreased vacuolar invertase activity (*amiR vi1-2*). Surprisingly, *BvTST2.1* overexpression led to increased monosaccharide levels in leaves, while sucrose remained constant. Latter observation allows the conclusion, that vacuolar invertase activity in mesophyll vacuoles exceeds sucrose uptake in Arabidopsis, which gained independent support by analyses on tobacco leaves transiently overexpressing *Bv*TST2.1 and the invertase inhibitor *Nb*VIF. However, we observed strongly increased sucrose levels in leaf extracts from independent *amiR vi1-2* lines and non-aqueous fractionations confirmed that sucrose accumulation in corresponding vacuoles. *amiR vi1-2* lines exhibited impaired early development and decreased weight of seeds. When germinated in the dark, mutant seedlings showed problems to convert sucrose into monosaccharides. Cold temperatures induced marked downregulation of the expression of both *VI* genes, while frost tolerance of *amiR vi1-2* mutants was similar to WT indicating that increased vacuolar sucrose levels fully compensate for low monosaccharide concentrations.

**Highlight:** Vacuolar sucrose accumulation in Arabidopsis is limited by high invertase activity and disturbed vacuolar sucrose homeostasis impairs plant germination, development, seed properties and survival under darkness.

## Introduction

Sugars are major solutes in plants and fulfil a plethora of functions in energy metabolism, synthesis of primary and secondary metabolites, lipid and protein modification, starch and cell wall biosynthesis, biotic and abiotic stress tolerance and intracellular signaling processes (Eveland and Jackson, 2011). In most species glucose and fructose, as typical monosaccharides, and the disaccharide sucrose represent major sugar types, although many other sugar varieties are also present in plants.

The importance of these three major types of sugars named above is not only given by the impressive number of cellular processes affected by them, but also by the observation that their individual cellular concentrations are sensed, and that corresponding information is translated into the expression of certain sets of genes (Koch, 2004; Koch, 1996; Cho and Yoo, 2011). This modification of gene expression couples cellular or developmental processes to the availability of different sugars. For example, high glucose levels are translated into a program initiating division and subsequent expansion of cells (Weschke *et al*., 2003), high fructose levels cause inhibition of seed development (Cho and Yoo, 2011) while high sucrose levels favor plant cell differentiation and maturation (Borisjuk *et al*., 2002; Koch, 2004).

In contrast to all other eukaryotes, plants generate sugars *de novo* during photosynthesis which usually starts with cytosolic sucrose biosynthesis in leaf mesophyll cells (Ruan, 2012). Subsequent to this, sucrose can be hydrolyzed by either, invertases or sucrose synthases, respectively, and the resulting reaction products serve e.g. as fuel for cellular energy provision via oxidative phosphorylation. Alternative to this, in most plant species, sucrose serves as the inter-organ, long distance transport form of carbohydrates (Lemoine *et al*., 2013). For latter purpose, sucrose is exported from the mesophyll cells, which is facilitated by sugar efflux transporters of the so called *Sugar Will Eventually be Transported* (SWEET) type (Chen, 2014). Subsequent to this, the phloem transport system is actively loaded with sucrose, which in case of Arabidopsis or sugar beet is driven by the proton-sucrose symporters belonging to the SUC2/SUT1 clade of sugar carriers (Truernit and Sauer, 1995; Gahrtz *et al*., 1994; Nieberl *et al*., 2017).

Therefore, sucrose in plant cells serves as a molecular signal governing gene expression, as a carbon storage molecule or as the main cargo to transport energy and carbon from source to sink organs. Although, sucrose biosynthesis occurs exclusively in the cytosol (Braun *et al*., 2014; Ruan, 2012) this sugar is also present in the apoplasm, and intracellularly in plastids and vacuoles (Heber, 1957; Klie *et al*., 2011; Hoermiller *et al*., 2017; Patzke *et al*., 2019). Therefore, it is not surprising that controlled hydrolysis of sucrose by either, cell wall-, vacuolar- or cytosolic invertases is critical for plant growth and properties (Heyer *et al*., 2004; Barratt *et al*., 2009; Wang *et al*., 2014; Weiszmann *et al*., 2018).

However, given the marked impact of sucrose for plant properties (Ruan, 2012) and having in mind that the vacuole represents the largest cell compartment (Martinoia *et al*., 2007) it is of special importance to characterize sucrose homeostasis in this organelle in more detail. Thus, to raise our knowledge on vacuolar sucrose homeostasis and its function for plant properties we generated various Arabidopsis mutants in which we either, ectopically expressed the sucrose specific vacuolar sugar loader TST2.1 from sugar beet (*Bv*TST2.1), or decreased the activity of vacuolar invertase (VI). With help of these mutants, we addressed following questions. 1. Does overexpression of *BvTST2.1* provoke sucrose storing in Arabidopsis leaves? 2. If not, what is the reason for this? 3. What is the impact of invertase-driven vacuolar sucrose hydrolysis for leaf sugar metabolism and plant properties? We provide evidences (i) for a substantial contribution of vacuolar invertase to a continuous sucrose turnover in Arabidopsis, (ii) that vacuolar sucrose import limits sucrose turnover, (iii) that decreased vacuolar sucrose turnover impairs early plant development and affects plant survival in phases of extended darkness and that (iv) vacuolar sucrose turnover is important for total seed yield. In sum of all analyses, we propose that a continuous vacuolar sucrose turnover is important for proper plant development.

## Materials and methods

### Plant material and growth conditions

*Nicotiana benthamiana*, *Arabidopsis thaliana* (ecotype Columbia-0) and corresponding Arabidopsis mutants were cultivated in a growth chamber (Weiss-Gallenkamp, Heidelberg, Gemany) on standardized soil (ED-73; Patzer; www.einheitserde.de), at a constant temperature of 22°C and a light intensity of 120 µmol quanta m^−2^ s^−1^ (µE). Plant cultivation was carried out under short day conditions,10 h light, 14 h darkness (standard conditions). For cold experiments, plants were grown for four weeks under standard conditions and subsequently acclimated to cold temperature for three days at 4°C (all other conditions were kept constant). For etiolation growth analyses, seeds were stratified for 24 h at 4°C, and cultivated in darkness for seven days on water soaked blotting paper. For seed analyses, plants were transferred after cultivation for four weeks at standard conditions to long day conditions (22°C and 200 µE, 16 h light per day). For dark recovery experiments, 4-week old plants grown under standard conditions were darkened for five days, followed by recovery for further 7 days under standard conditions.

### Generation of mutants

For transient transformation of *Nicotiana benthamiana* leaf mesophyll cells the Agrobacterium infiltration method was performed according to an established method (Jung *et al*., 2015). For generation of *VI1-2* double knock down mutants, we followed an established protocol for gene silencing by artificial microRNA (amiRNA) (Schwab *et al*., 2006). With the web based amiRNA designer tool program (http://wmd3.weigelworld.org) an amiRNA, which targets *VI1* (*At1g62660*) and *VI2* (*At1g12240*) simultaneously, was designed. The sequence *TAAGGATGAATAAAAGCACGG* was used for generation of primers including the Gateway™ compatible sequences attP1 and attP2 to engineer the amiRNA fragment. The primer sequences are listed in Table S1. Subsequently, the fragment was sub-cloned via BP reaction into the Gateway™ entry vector pDONR/Zeo and via LR reaction into the destination vector pK2GW7, which contains a 35S-CaMV promotor. For generation of stably transformed Arabidopsis mutant plants, Agrobacterium-mediated transformation using floral dip was performed (Clough and Bent, 1998). *VI1-2* double knock-down mutants were selected by screening for the lowest remaining *VI1* and *VI2* transcript levels via qRT-PCR leading to the two independent lines no. 4 and 5.

### cDNA synthesis, qRT-PCR and RNA gel-blot hybridization

RNA was isolated from 50 mg of frozen, fine ground plant material with the NucleoSpin RNA Plant Kit (Macherey-Nagel, Düren, Germany), according to the manufacturer’s protocol. For cDNA synthesis, RNA was transcribed into cDNA with the qScript cDNA Synthesis Kit (Quantabio, Beverly, MA, USA). The primers used for gene expression analysis by qRT-PCR are listed in Table S2. *NbAct*, *AtPP2A* and *AtSAND* were used as reference genes for transcript normalization. Alternatively, gene expression was analyzed by RNA gel-blot hybridization.

### Acidic invertase activity assay

The enzyme assay was performed as described by (Tamoi *et al*., 2010) with slight modifications. 100 mg of frozen and fine ground plant material were homogenized with 1 ml of ice cold 200 mM Hepes/HCl (pH 5.0), 1 mM EDTA, 1 mM PMSF on a vortex mixer for 20 sec. The samples were incubated for 20 min on ice prior to further mixing with a vortex mixer for 20 sec. Subsequently, samples were centrifuged at 20.000 *g* for 10 min at 4°C and the supernatant was transferred into a new reaction tube. The rate of sucrose hydrolysis was quantified spectrophotometrically at 22°C using a NADP-coupled enzymatic test (Stitt *et al*., 1989). For this, 15 µl of enzyme extract were added to 190 µl 200 mM Hepes/HCl (pH 5.0), 10.5 mM MgCl_2_, 2.1 mM ATP, 0.8 mM NADP, 0.5 U Glucose-6-Phosphate Dehydrogenase, 0.18 U Hexokinase and 0.48 U Phosphoglucose Isomerase. To start the enzyme reaction, 5 µl of 200 mM sucrose solution were added to the sample.

### Metabolite quantification

For sugar extraction, we added 400 µl of 80% of ethanol to 100 mg of frozen, fine grounded plant material, mixed and incubated for 30 min at 80°C, in a thermomixer at 500 rpm. After centrifugation at 16000 *g* (10 min at 4°C) the supernatant was used for sugar quantification using a NADP-coupled enzymatic test (Stitt *et al*., 1989).

### Non-aqueous fractionation

Subcellular sugar distribution of sugars in leaves was determined by non-aqueous fractionation of 4-week old plants. To this end, 15 mg of freeze-dried and fine grounded plant material was used and processed as described previously (Fürtauer *et al*., 2016). Acid phosphatase served as vacuolar marker, UGPase activity served as a cytosolic marker and alkaline pyrophosphatase served as chloroplast marker. For sugar quantification, a NADP-coupled enzymatic test was performed (Stitt *et al*., 1989) and subcellular metabolite distribution was calculated using an established algorithm (Fürtauer *et al*., 2016).

### Seed analyses

Lipid quantification was performed according to a routine protocol (Reiser *et al*., 2004) with slight modifications. 0.1 g of mature, air-dried seeds were homogenized in a mortar and liquid nitrogen. Subsequently, 1.5 mL of isopropanol was added and the sample was further homogenized. The suspension was transferred into a 1.5 mL-reaction tube and incubated for 12 h at 4°C and 100 rpm. Subsequently, samples were centrifuged at 13000 *g* for 10 min and the supernatant was transferred into a pre-weighed 1.5-mL reaction tube. Tubes were incubated at 60°C for 12 h to completely evaporate the isopropanol. Total lipid content was quantified gravimetrically. For determination of seed weight, 1000 mature and air-dried seeds were counted and their weight was quantified gravimetrically.

### Electrical conductivity

The frost tolerance, as measured by frost induced release of ions from leaf sample, of wild-type and mutant, was quantified by electrical conductivity assays, described earlier (Klemens *et al*., 2014).

## Results

### Overexpression of either, the vacuolar sucrose loader TST2.1 or the vacuolar sucrose exporter SUC4 provoke opposite effects on cellular sugar levels

The plant vacuole is the largest cell organelle and as such uniquely suited to store various types of sugars (Martinoia *et al*., 2012). Given that *Beta vulgaris* TST2.1 is a sucrose specific vacuolar sugar loader (Jung *et al*., 2015) we tried to stimulate leaf sucrose storage capacity by overexpression of this carrier. To this end, we set the coding sequence of the *BvTST2.1* gene under control of the *UBIQUITIN10*-promotor (*UBQ-10*) and transformed a previously described *Arabidopsis thaliana tst1-2* double mutant, lacking almost all vacuolar sugar import (Wingenter *et al*., 2010). The resulting triple mutants were named *tst1-2::BvTST2.1*.

As expected, neither in wild-type (WT) nor in the *tst1-2* line, used as the genetic basis for transformation, a PCR product coding for *Bv*TST2.1 was detectable (Fig. S1). In contrast, in transformed *tst1-2* plants, high *BvTST2.1* mRNA levels were present. From about 25 independent overexpressing lines two were chosen (*tst1-2::BvTST2.1* #18 and *tst1-2::BvTST2.1* #23) for further analyses because they exhibited high relative levels of *BvTST2.1* transcripts (Fig. S1).

To analyze whether this synthetic approach sufficed to create a novel sucrose storing organ we grew all plants for 28 days under controlled, ambient cultivation conditions and extracted leaf metabolites at the end of the light phase, prior to quantification of sugars (Fig. 1A). In WT plants the levels of glucose, fructose and sucrose were 2.2 µmol g^−1^ FW, 0.8 µmol g^−1^ FW and 1.3 µmol g^−1^ FW, respectively (Fig. 1A). In contrast, *tst1-2* mutants were unable to build up such monosaccharide concentrations in leaves as glucose was present at only 0.9 µmol g^−1^ FW and fructose was present at about 0.4 µmol g^−1^ FW. In the *tst1-2* mutant sucrose accumulated to 1.5 µmol g^−1^ FW, which is about the similar level found in corresponding WT plants (Fig. 1A). Interestingly, overexpression of *BvTST2.1*, in the background of *tst1-2*, led to higher glucose (3.1 and 2.6 µmol g^−1^ FW in line 18 and 23) and fructose (1.2 and 0.9 µmol g^−1^ FW in line #18 and #23) levels, and thus resembled corresponding contents present in WT plants. In contrast, sucrose levels in *BvTST2.1* overexpressing lines #18 and #23 amounted to 1.1 and 1.3 µmol g^−1^ FW, respectively, which were slightly (but not yet significantly) decreased compared to values in the *tst1-2* background (Fig. 1A).

**Fig. 1.**
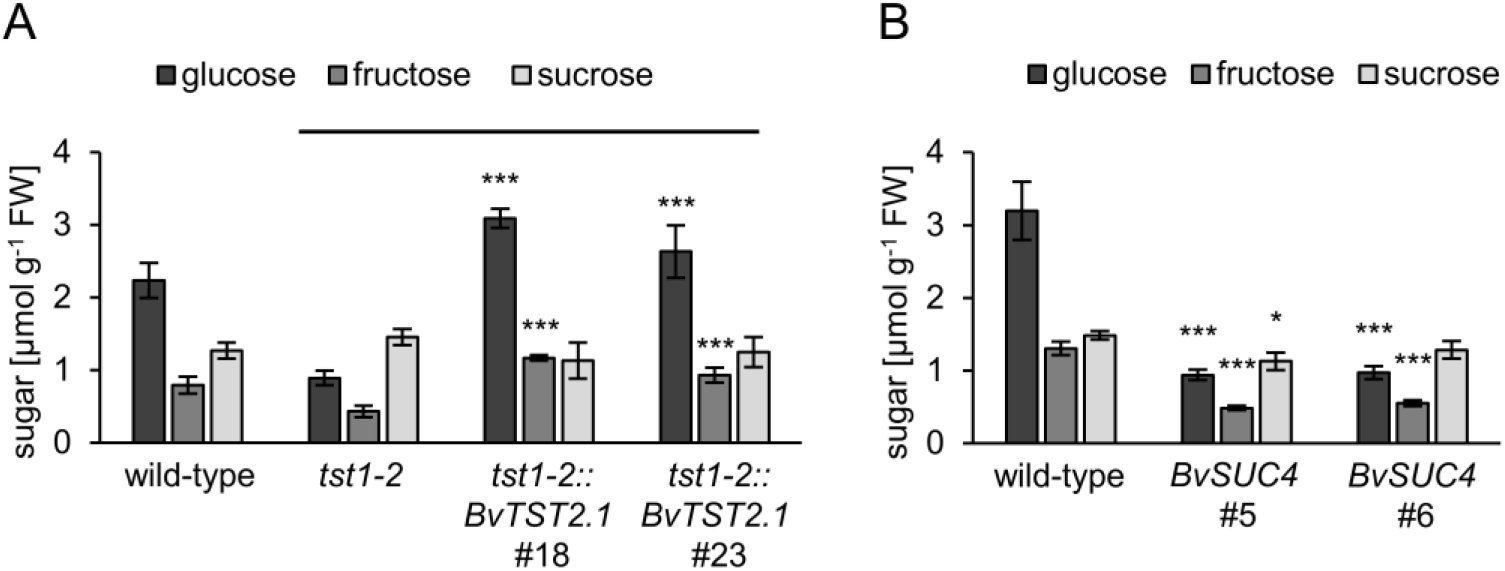
Soluble sugar contents of 4-week old plants grown on soil. **A** Sugar levels of *tst1-2* mutant, *tst1-2::BvTST2.1* overexpressor line 18 and 23 and the corresponding wild-type. Data are presented as mean ±SE of at least 6 biological replicates. **B** Sugar levels of *BvSUC4* overexpressor line 5 and 6 and the corresponding wild-type. Data are presented as mean ±SE of at least 4 biological replicates. Asterisks indicate statistically significant differences between the *tst1-2* and the *tst1-2::BvTST2.1* lines or between the *BvSUC4* lines and the corresponding wild-type analyzed with Students t-test (* p≤0.05; *** P≤0.001).

To raise further evidence that altered sucrose transport across the vacuolar membrane influences the cellular sucrose to monosaccharide ratio we searched for an opposite attempt aiming to decrease the luminal sucrose availability. For this, we identified the sugar beet homolog of the vacuolar sucrose transporter *At*SUC4. *At*SUC4 is a proton driven sucrose exporter (Schneider *et al*., 2012) and in sugar beet (*Beta vulgaris*) the carrier *Bv*SUC4 represents the closest homolog to *At*SUC4 (Dohm *et al*., 2014), exhibiting 67% identity of the amino acid sequence (BLASTP; https://blast.ncbi.nlm.nih.gov/Blast.cgi). After transformation of Arabidopsis plants with a *35S::BvSUC4* construct, lines *BvSUC4 #5* and *BvSUC4 #6* have been identified as overexpressors, exhibiting high levels with corresponding ectopic mRNAs (Fig. S2A,B).

After growth for four weeks at standard conditions WT plants contained about 3.2 µmol glucose g^−1^ FW, 1.3 µmol fructose g^−1^ FW and 1.5 µmol sucrose g^−1^ FW (Fig. 1B). In contrast, both *BvSUC4* overexpressing lines contained less monosaccharides, while sucrose levels were similar to WT levels. *BvSUC4* #5 contained 0.9 µmol glucose g^−1^ FW, 0.5 µmol fructose g^−1^ FW and 1.1 µmol sucrose g^−1^ FW, while and *BvSUC4* #6 contained 1.0 µmol glucose g^−1^ FW, 0.6 µmol fructose g^−1^ FW and 1.3 µmol sucrose g^−1^ FW (Fig. 1B).

### Vacuolar invertase contributes to intracellular sucrose turnover

The metabolite data above indicate that altered sucrose transport across the vacuolar membrane influenced the cellular sucrose to monosaccharide ratio. Moreover, overexpression of *BvTST2.1* does not allow the conversion of Arabidopsis leaves into high sucrose storing organelles. One explanation for these observations is that sucrose hydrolysis in Arabidopsis vacuoles from *BvTST2.1* overexpressing lines is faster than in corresponding WT plants. To raise independent experimental evidence for an existence of such efficient sucrose hydrolysis in mesophyll vacuoles via invertase, we exploited a gene expression system based on transient transformation of tobacco leaf mesophyll cells with help of *Agrobacterium tumefaciens* (Zhao *et al*., 2017).

For this purpose, we transiently transformed tobacco mesophyll cells with Agrobacterium strains carrying plasmids harbouring the viral silencing suppressor gene P19 from *tomato bushy stunt virus* (Voinnet *et al*., 2003) alone or in combination with either (i) the vacuolar invertase inhibitor from *Nicotiana benthamiana* (P19/*NbVIF*), (ii) the *BvTST2.1* gene (P19/*BvTST2.1*), or (iii) *NbVIF* and *BvTST2.1 (*P19/*BvTST2.1*/*NbVIF)* (Fig. 2A). Prior to metabolite analyses, we confirmed that transient transformation leads always to the presence of the expected corresponding mRNA(s) (Fig. 2B).

**Fig. 2.**
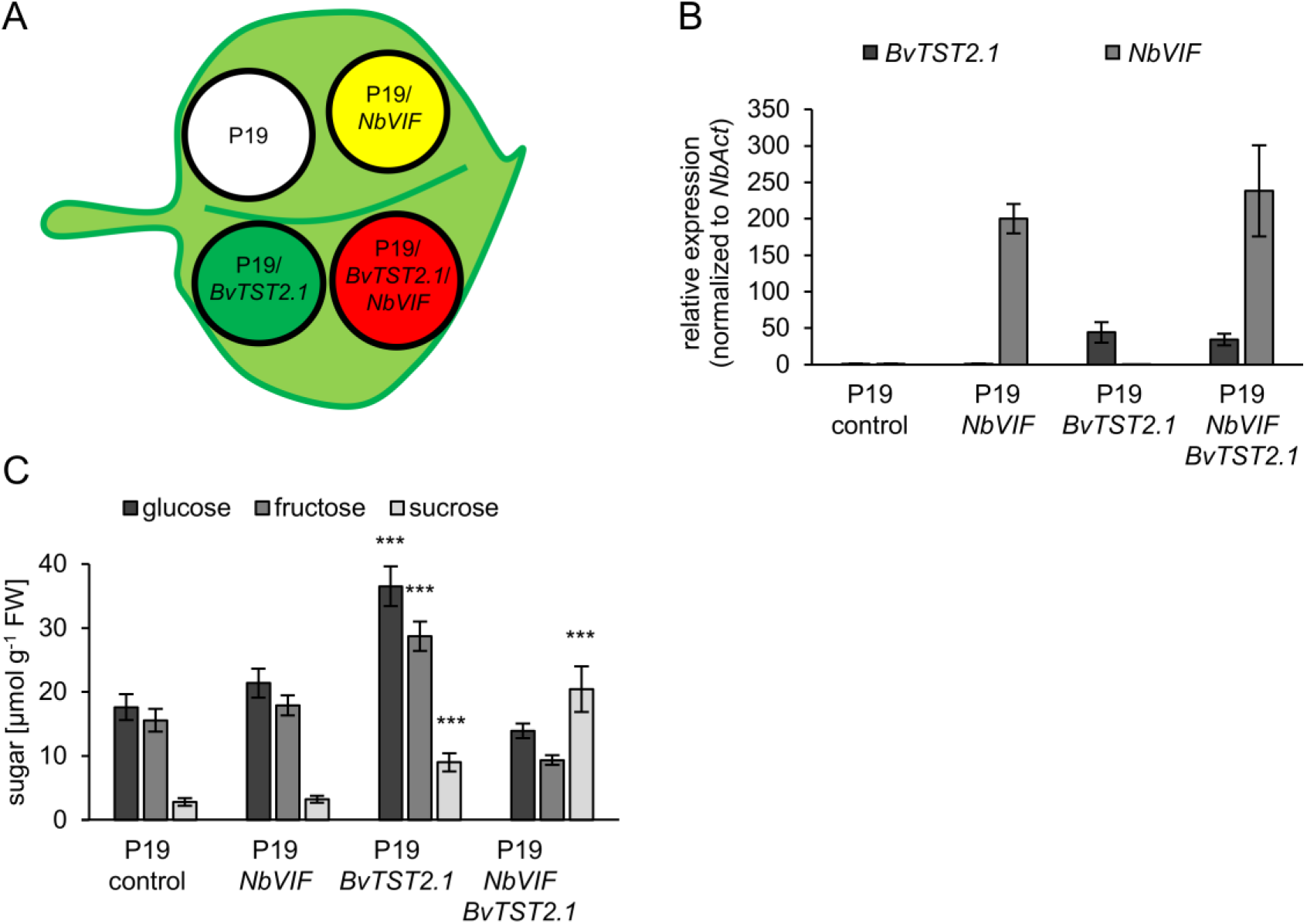
Elucidation of *Bv*TST2.1’s *in vivo* function using *N. benthamiana* infiltration assay. **A** Schematic drawing of a *N. benthamiana* leaf infiltrated with Agrobacteria harboring different expression constructs. **B** Normalized expression of *BvTST2.1* and *NbVIF* in infiltrated leaf tissue area. **C** Soluble sugar levels of leaf tissue harvested 4 days after infiltration. Data are presented as mean ±SE of at least 6 biological replicates. Asterisks indicate statistically significant differences analyzed with Students t-test (*** P≤0.001). P19 = P19 protein of *tomato bushy stunt virus*, a suppressor of gene silencing (Voinnet *et al*., 2003); *NbVIF* = inhibitor protein of *N. benthamiana* vacuolar invertase.

Infiltrated leaves were left attached to plants, which were further cultivated for 48 h under the indicated growth conditions until extraction of metabolites from the four different infiltrated leaf areas (at the end of the light phase). In the control area (P19) glucose and fructose were present at a concentration of about 17 µmol g^−1^ FW, while sucrose was present at 3 µmol g^−1^ FW (Fig. 2C). In leaf tissue infiltrated with the plasmid P19/*NbVIF* the levels of monosaccharides and sucrose were similar to the leaf area infiltrated with the control plasmid P19 (Fig. 2C). Leaf tissue expressing *BvTST2.1* (P19/*BvTST2.1*) substantially increased levels of glucose, fructose and sucrose were present, when compared to the control area P19 (Fig. 2C). Remarkably, the simultaneous expression of the vacuolar invertase inhibitor *NbVIF* and the carrier *BvTST2.1* led to a greatly increased accumulation of sucrose, while levels of glucose and fructose were similar in comparison to control-infiltrated P19 tissue (Fig. 2C).

### An induced decrease of vacuolar invertase activity affects total sucrose levels and modifies intracellular sucrose compartmentation in Arabidopsis leaves

In Arabidopsis, two acidic, vacuolar located invertases have been annotated namely *At1g62660* (*VI1*) and *At1g12240* (*VI2*). Recently it has been shown, that lack of a functional *VI2* gene, coding for the isoform contributing most of the total vacuolar invertase activity, does not provoke significantly altered vacuolar sugar levels under both, ambient temperature or exposure to cold temperature (Weiszmann *et al*., 2018a). Thus, to create mutant plants with decreased activities of both vacuolar invertases, we created, on basis of Arabidopsis WT, artificial microRNA mutants exhibiting strongly decreased levels of mRNA coding for both vacuolar invertases (*amiR vi1-2*) (Fig. 3). From 10 independent mutant lines we chose lines #4 and #5 for further analysis, because with 6 to about 24% residual *VI1* and *VI2* mRNA levels, respectively, these two lines showed strongest decrease of the corresponding transcripts (Fig. 3A). Consequently, measurement of acidic invertase activities in *amiR vi1-2* #4 and #5 revealed only 22 and 35% residual enzyme activity, respectively (Fig. 3B).

**Fig. 3.**
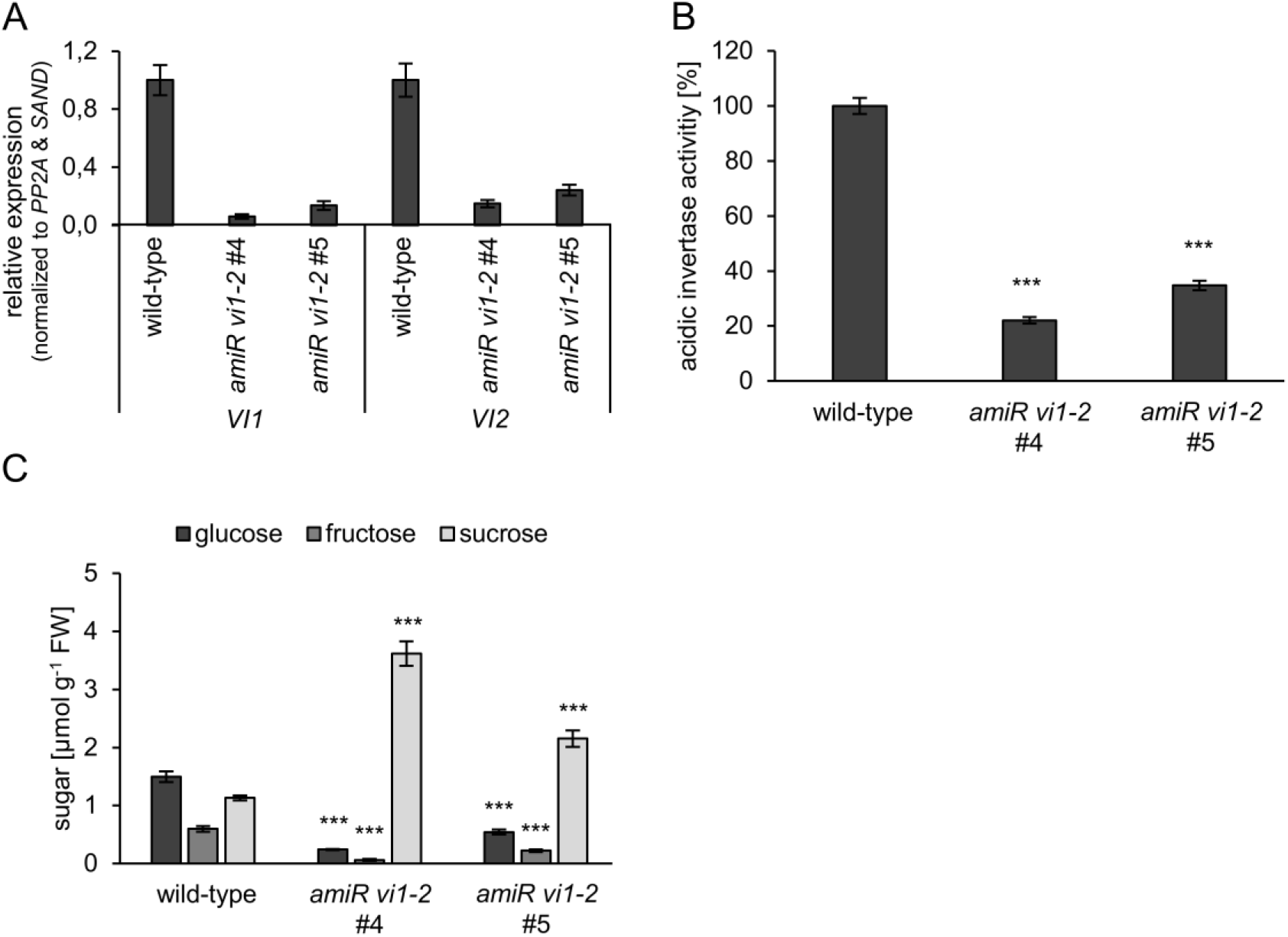
Molecular, biochemical characterization and soluble sugar content of 4-week old wild-type plants and *amiR vi1-2* lines 4 and 5. **A** Normalized relative expression level of *vacuolar invertase* (*VI*) *1* and *2* in leaf samples Data are presented as mean ±SE of 4 biological replicates. **B** Acidic invertase activity in leaf samples. Data are presented as mean ±SE of 5 biological replicates. **C** Sugar levels in leaves of 4-week old plants grown on soil. Data are presented as mean ±SE of 6 biological replicates. Asterisks indicate statistically significant differences between the wild-type and the *amiR vi1-2* lines analyzed with Students t-test (*** P≤0.001).

When grown under standard temperature (22°C), WT plants contained, after 4 weeks of cultivation, glucose at a level of 1.5 µmol g^−1^ FW, fructose at a level of 0.6 µmol g^−1^ FW and sucrose at a level of 1.1 µmol g^−1^ FW (Fig. 3C). Interestingly, the *amiR vi1-2* lines exhibited less monosaccharides and increased sucrose levels when compared to corresponding WT plants. Mutant line #4 contained glucose at a level of only 0.2 µmol g^−1^ FW, fructose at a level of only 0.04 µmol g^−1^ FW and sucrose at a level of 3.6 µmol g^−1^ FW (Fig. 3C). Mutant line #5 contained glucose at a level of only 0.5 µmol g^−1^ FW, fructose at a level of only 0.2 µmol g^−1^ FW and sucrose at a level of 2.2 µmol g^−1^ FW (Fig. 3C).

To analyse whether a decrease of vacuolar invertase activity would influence not only the overall sucrose levels but also the intracellular sucrose compartmentation in Arabidopsis leaves, we conducted non-aqueous fractionation, allowing to determine the relative distribution of sugars between chloroplasts, cytosol and vacuoles (Fürtauer *et al*., 2016). Although both *amiR vi1-2* lines exhibited some changes in the intracellular distribution of glucose and fructose, when compared to WT plants (Fig. 4A,B), the most prominent changes were observed for sucrose concentrations in the three cell compartments analysed (Fig. 4C). WT stored about 18% of the cellular sucrose in the vacuole, while *amiR vi1-2 #4* plants stored about 58% and *amiR vi1-2 #5* plants about 38% of the total cell sucrose in the vacuole (Fig. 4C). Interestingly, increased vacuolar sucrose levels in both *amiR vi1-2* lines correlated with decreased sucrose levels in corresponding chloroplasts (Fig. 4C), known as another cellular site where sucrose occurs in mono- and dicotyledonous plants (Heber, 1957; Gerhardt *et al*., 1987).

**Fig. 4.**
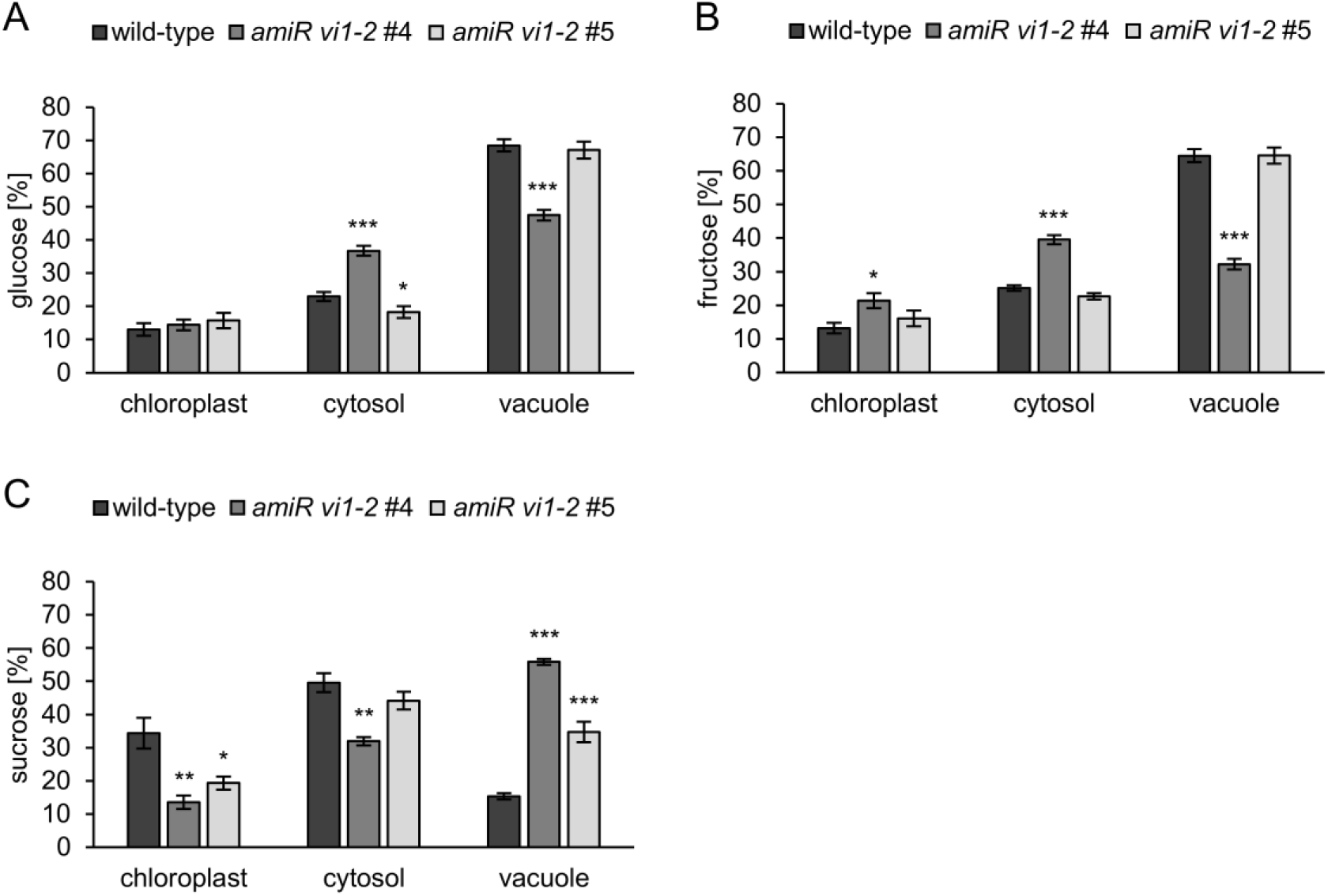
Analysis of subcellular sugar distribution of 4-week old wild-type plants and *amiR vi1-2* lines 4 and 5 after non-aqueous fractionation. Subcellular distribution of glucose (**A**), fructose (**B**) and sucrose (**C**). Data are presented as mean ±SE of 4 biological replicates consisting of 3 plants each. Asterisks indicate statistically significant differences between the wild-type and the *amiR vi1-2* lines analyzed with Students t-test (* P≤0.05; ** P≤0.01; *** P≤0.001).

### Reduced vacuolar invertase activity impacts efficient early seedling development under dark growth conditions

Seeds from most plants are able to germinate allowing growth in the absence of light, leading to an etiolated habitus. Under these conditions, seedlings (e.g. from Arabidopsis) are fully dependent upon an efficient conversion of storage lipids to sugars. However, it is unknown whether vacuolar invertase activity influences this metabolic phenomenon allowing etiolated growth, termed gluconeogenesis.

Thus, to provide insight into a putative function of vacuolar invertase activity for plant growth under dark conditions seeds from WT and *amiR vi1-2* were stratified and germinated seedlings cultivated for 7 days in darkness. Subsequently, we determined shoot length and sugar levels of these seedlings. Both *amiR vi1-2* lines exhibited decreased length of the seedlings when compared to simultaneously grown WT seedlings (Fig. 5A). In average, etiolated WT seedlings exhibited a length of 15 mm, while seedlings from *amiR vi1-2* #4 exhibited only 13 mm and seedlings from *amiR vi1-2* #5 exhibited only 12 mm (Fig. 5B). Interestingly, WT seedlings contained about 7.9 µmol glucose g^−1^ FW, about 2.4 µmol fructose g^−1^ FW and 0.7 µmol sucrose g^−1^ FW. In contrast, *amiR vi1-2* #4 contained 1.5 µmol glucose g^−1^ FW, 0.1 µmol fructose g^−1^ FW, and 2.7 µmol sucrose g^−1^ FW. *AmiR vi1-2* #5 contained (similar to *#*4) 1.0 µmol glucose g^−1^ FW, 0.06 µmol fructose g^−1^ FW and 1.8 µmol sucrose g^−1^ FW (Fig. 5C). In sum, WT seedlings contained about 11.6 µmol hexoses g^−1^ FW, while both *amiR vi1-2* lines contained only 7 and 5 µmol hexoses g^−1^ FW, respectively (Fig. 5D).

**Fig. 5.**
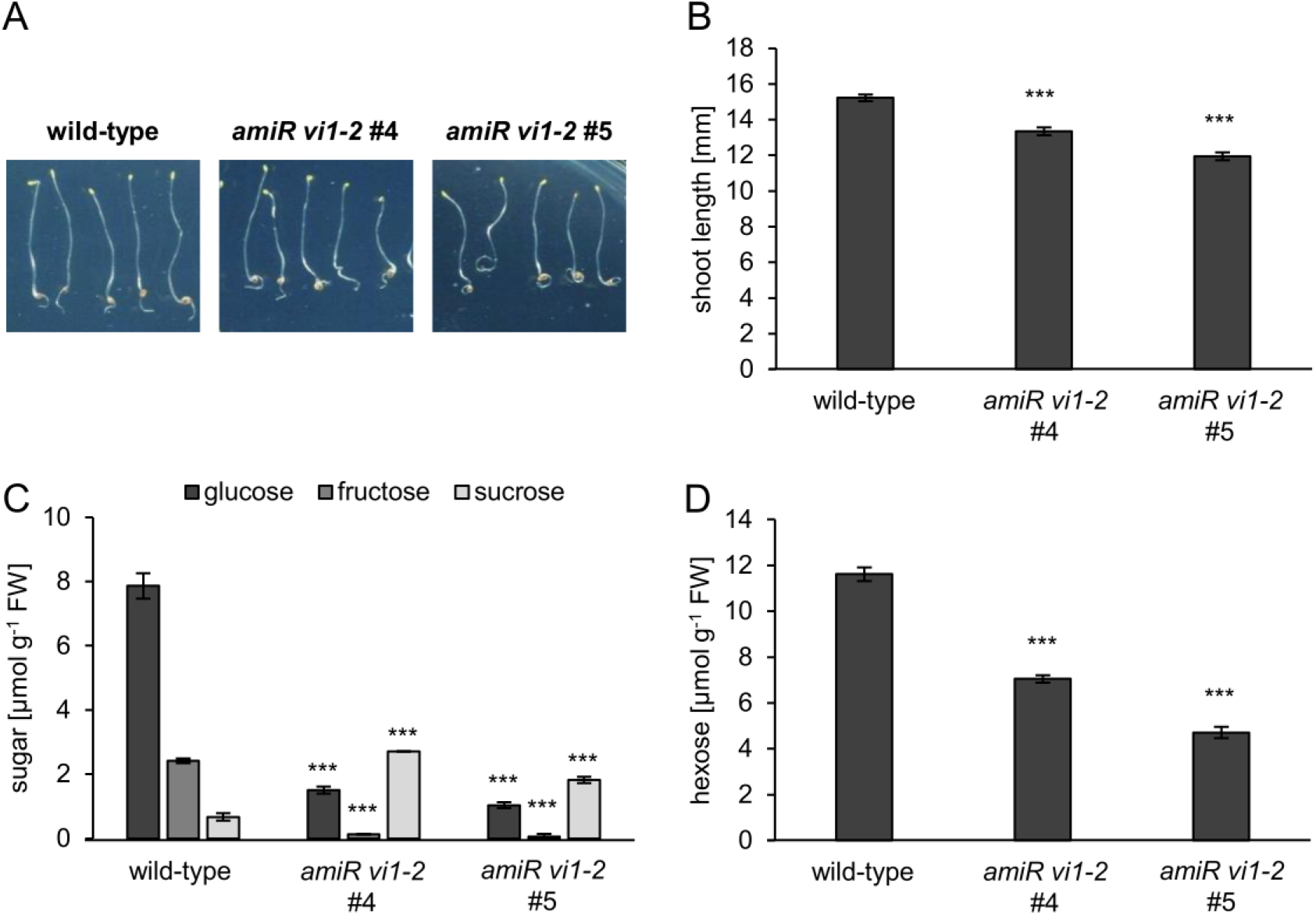
Effects of dark treatment on plant phenotype and sugar levels of wild-type plants and *amiR vi1-2* lines 4 and 5 after germination for 7 days in darkness. **A** Etiolated seedlings. **B** Analysis of etiolated shoot lengths. Data are presented as mean ±SE of at least 30 biological replicates. **C** Sugar levels in etiolated seedlings. Data are presented as mean ±SE of 3 biological replicates. **D** Total hexose levels in etiolated seedlings. Data are presented as mean ±SE of 3 biological replicates. Asterisks indicate statistically significant differences between the wild-type and the *amiR vi1-2* lines analyzed with Students t-test (*** P≤0.001).

### Reduced vacuolar invertase activity impacts on seed properties

It is unknown whether decreased vacuolar invertase affects seed properties. To raise evidence for an interaction between VI activity and seed properties, we grew WT and both *amiR vi1-2* lines for five weeks under short day conditions (10 h light phase) and transferred all plants subsequently to long day (16 h light phase) conditions to induce flowering. Seeds were harvested after plants completed their life cycle, and weight and lipid content of dry seeds were quantified. In addition, we quantified total sugar levels in developed siliques.

To exclude that putative changes in seed properties result from differences in the vegetative parts of the plants we first determined the biomasses of all three plant lines prior to transfer to long days. It turned out, that all plant lines analysed, WT and the two *amiR vi1-2* lines, exhibited nearly identical rosette biomasses of about 730 mg/plant (Fig. 6A,B). The 1000-seed weight from WT plants was 25 mg, and those from the *amiR vi1-2* #4 and #5 was 23 mg and 20mg, respectively (Fig. 6C). Accordingly, the relative lipid levels of seeds from both *amiR vi1-2* lines were slightly, but still significantly, lower than that of seeds from WT plants. *amiR vi1-2* #4 seeds contained 89% and *amiR vi1-2* #5 seeds 93% of WT lipids (Fig. 6D).

**Fig. 6.**
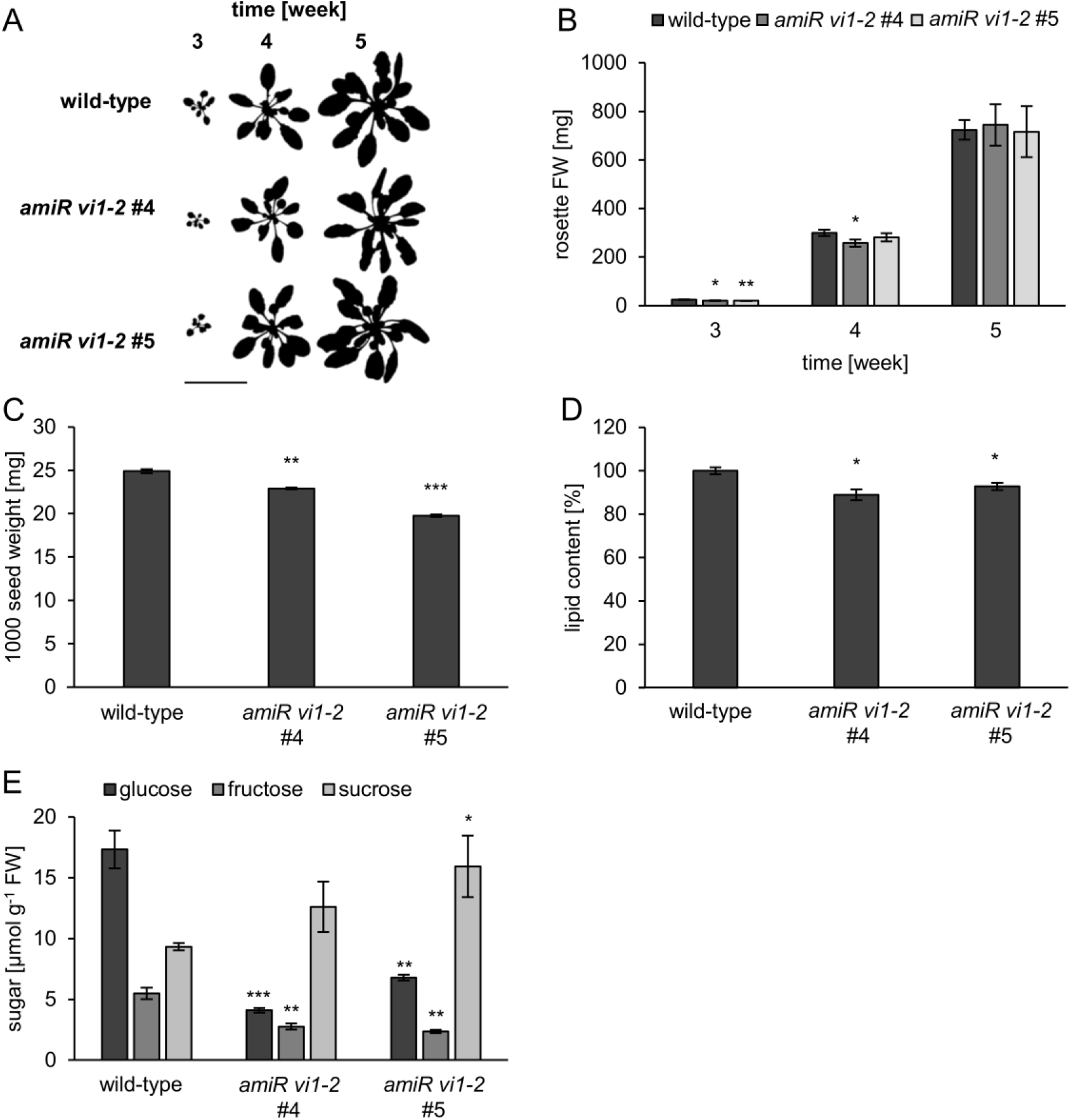
Analysis of rosettes, seeds and siliques of wild-type plants and *amiR vi1-2* lines 4 and 5. **A** Rosette size of 3- to 5-week old plants. Bar = 5 cm. **B** Analysis of rosette fresh weight of 3- to 5-week old plants. Data are presented as mean ±SE of at least 6 biological replicates. **C** 1000 seed weight. Data are presented as mean ±SE of 3 biological replicates (deriving from the same harvest). **D** Lipid content of mutant seeds was normalized to lipid content of wild-type seeds. Data are presented as mean ±SE of 4 biological replicates. **E** Sugar content in siliques. Data are presented as mean ±SE of 3 biological replicates. Asterisks indicate statistically significant differences between the wild-type and the *amiR vi1-2* lines analyzed with Students t-test (* P≤0.05; ** P≤0.01; *** P≤0.001).

To check whether altered VI activity in both *amiR vi1-2* lines influences the overall sugar composition in siliques in which seeds were developing, we harvested fully elongated, but still green siliques and extracted soluble carbohydrates. WT siliques contained about 17 µmol glucose g^−1^ FW, 5 µmol fructose g^−1^ FW, and 9 µmol sucrose g^−1^ FW, while siliques harvested from *amiR vi1-2* #4 contained only 4 µmol glucose g^−1^ FW, 3 µmol fructose g^−1^ FW but 13 µmol sucrose g^−1^ FW and siliques harvested from *amiR vi1-2* #5 contained only 7 µmol glucose g^−1^ FW, 2 µmol fructose g^−1^ FW but 16 µmol sucrose g^−1^ FW (Fig. 6E).

### Vacuolar invertase activity influences the endurance of Arabidopsis plants under continuous dark conditions

It appears likely that under conditions of a low energy status, vacuolar invertases contribute to supply the cell with carbohydrates required for oxidative phosphorylation. To verify this hypothesis, we grew incubated Arabidopsis WT and the two *amiR vi1-2* lines for 4 weeks under standard short day conditions prior to darkening of the plants for up to five days.

To raise experimental evidence for a putative involvement of the vacuolar invertase we first tested whether the expression of either the *VI1* or *VI2* gene responded to extended dark period. It turned out, that the level of *VI1* mRNA was not altered in leaves upon onset of extended darkness, while *VI2* mRNA increased up to 3-fold within 24 h of darkness (Fig. S3A). Concomitantly, extractable acidic invertase activity from corresponding dark incubated leaves increased up to 280% of the value at the beginning of the dark treatment (Fig. S3B).

To visualize and quantify the ability of WT and the *amiR vi1-2* lines to survive extended darkness we incubated plants from these lines for five days under continuous darkness and subsequently transferred them back into the standard short day light/dark regime for recovering. After one additional week, it became obvious, that both *amiR vi1-2* lines showed higher tolerance against this extended darkness in comparison to WT plants (Fig. 7A). About 57% of the WT plants were able to recover from the extended darkness treatment, while 82% of *amiR vi1-2* #4 plants and 75% of the *amiR vi1-2* #5 plants were able to survive such phase of darkness (Fig. 7B).

**Fig. 7.**
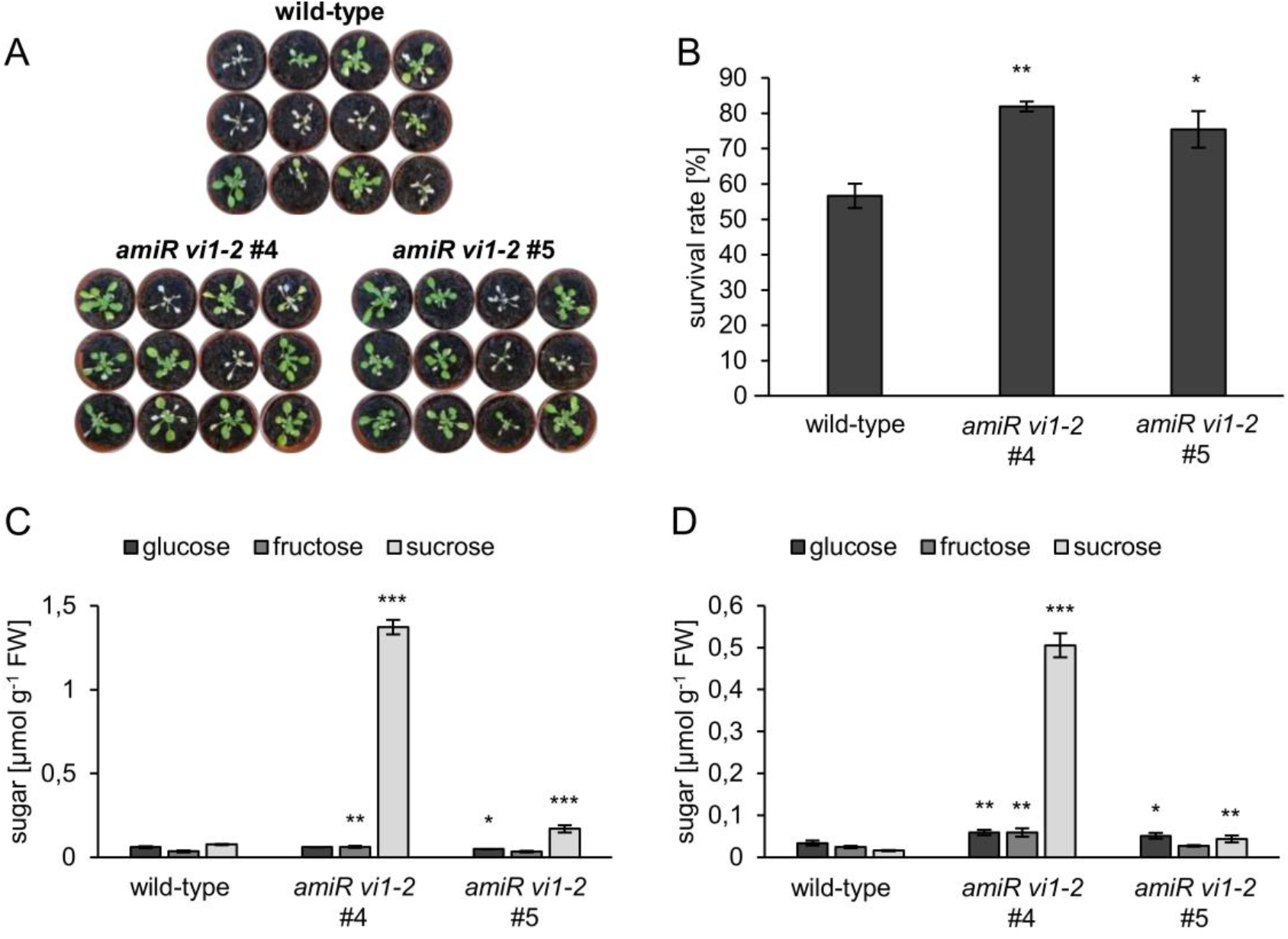
Effects of dark treatment on wild-type plants and *amiR vi1-2* lines 4 and 5. Plants were cultivated for 4 weeks under standard conditions on soil, kept for 5 days in the dark and then recovered for 7 days under standard conditions. **A** Plants after dark recovery. **B** Quantification of survivors after dark recovery. Data are presented as mean ±SE of 3 independent experiments with each 12 plants per line. Sugar levels in leaves after 24 hours (**C**) and after 72 hours (**D**) of dark treatment. Data are presented as mean ±SE of 4 biological replicates. Asterisks indicate statistically significant differences between the wild-type and the *amiR vi1-2* lines analyzed with Students t-test (* P≤0.05; ** P≤0.01; *** P≤0.001).

To raise knowledge on specific metabolic alterations in these plant lines we extracted sugars from plants after 24 and 72 h of darkness. After 24 h of darkness, WT plants exhibited markedly low levels of all tree sugars, since each glucose, fructose and sucrose ranged only between 0.06 and 0.08 µmol g^−1^ FW (Fig. 7C). In contrast, both *amiR vi1-2* lines showed higher levels of sucrose in comparison to WT plants. Leaves from *amiR vi1-2* #4 plants contained 1.37 µmol sucrose g^−1^ FW, and *amiR vi1-2* #5 leaves contained two times higher sucrose levels than present in corresponding WT plants, namely 0.17 µmol sucrose g^−1^ FW (Fig. 7C). Interestingly, even after 72 h of darkness both *amiR vi1-2* lines contained still higher sugar levels than present in WT plants. Leaves from *amiR vi1-2* #4 plants contained about 30-fold higher sucrose levels than present in WT plants, and about 2-fold higher monosaccharide concentrations and *amiR vi1-2* #5 leaves contained 3-fold higher sucrose levels and nearly 2-fold higher glucose levels than present in WT plants (Fig. 7D).

### *AmiR vi1-2* lines showed marked alterations of intracellular sugar levels in the cold but no modification of frost tolerance

Intracellular sugar homeostasis is of marked importance for plant frost tolerance (Nägele *et al*., 2010; Nägele *et al*., 2012; Pommerrenig *et al*., 2018). To raise further evidence on the putative function of vacuolar sucrose levels we first analysed the response of *VI1* and *VI2* gene expression upon onset of cold temperatures. After three days at 4°C, both mRNAs dropped significantly and reached only between 12 and 10% of the mRNA values, respectively, present in plants grown at 22°C (Fig. 8A).

**Fig. 8.**
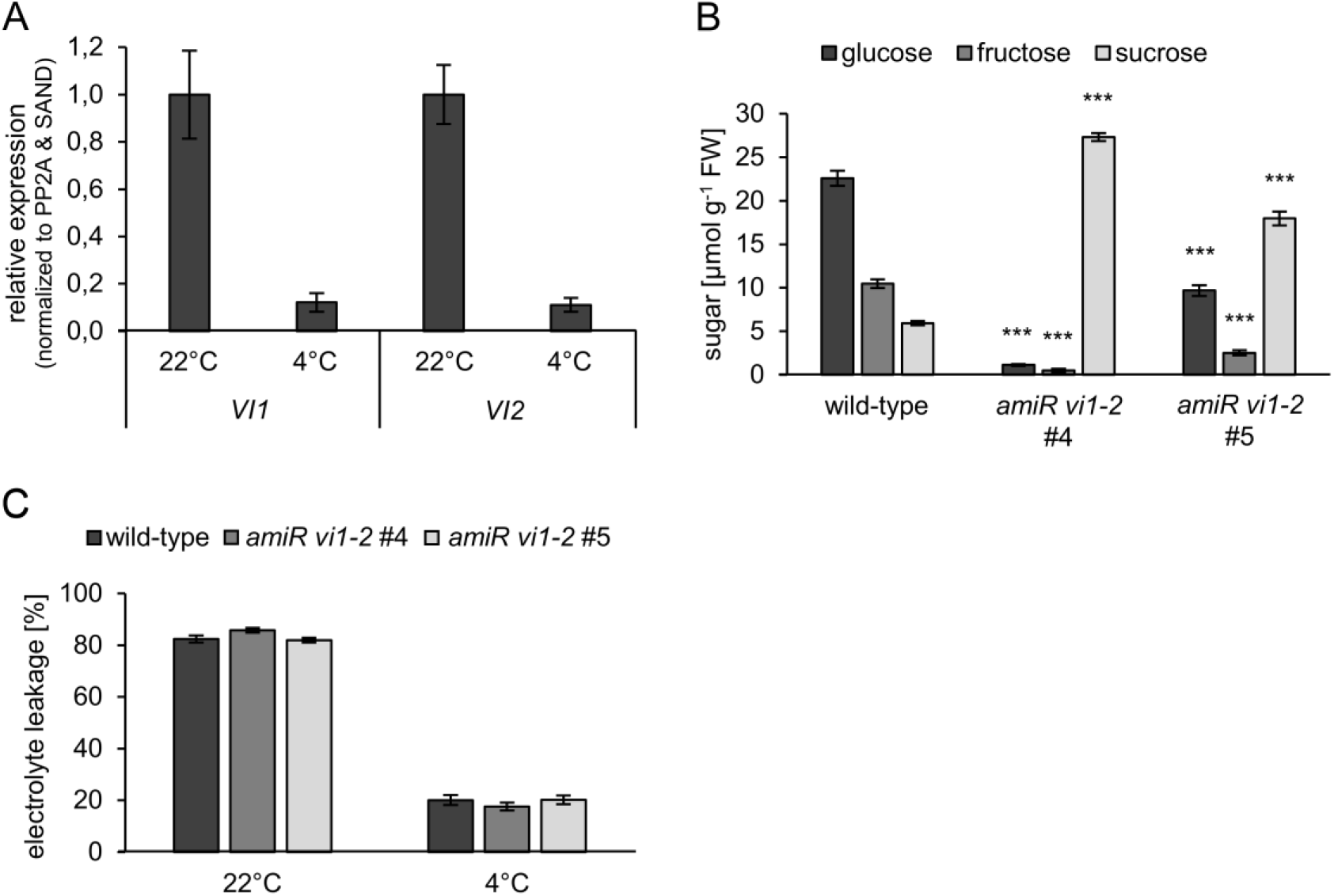
Effects of cold treatment on wild-type plants and *amiR vi1-2* lines 4 and 5. Plants were cultivated for 4 weeks under standard conditions on soil and then transferred for 3 days to 4°C. **A** Relative expression level of *vacuolar invertase* (*VI*) *1* and *2* in cold acclimated wild-type leaf samples. Data are presented as mean ±SE of 4 biological replicates. **B** Sugar levels in cold acclimated leaves. Data are presented as mean ±SE of 6 biological replicates. **C** Analysis of electrolyte leakage of leaves kept in cold (4°C) for 4 days. Data are presented as mean ±SE of at least 8 biological replicates. Asterisks indicate statistically significant differences between the wild-type and the *amiR vi1-2* lines analyzed with Students t-test (*** P≤0.001).

To compare the cold induced sugar response in WT and the two *amiR vi1-2* lines, we cultivated 4-week old plants (grown at 22°C) for three days at 4°C. After that acclimation phase, WT plants contained glucose at a level of 23 µmol g^−1^ FW, fructose at a level of 10 µmol g^−1^ FW and sucrose at a level of 6 µmol g^−1^ FW (Fig. 8B). As observed under warm growth conditions, the two *amiR vi1-2* lines exhibited less accumulation of monosaccharides but increased accumulation of sucrose, when compared to corresponding WT plants. In the cold, mutant line #4 contained glucose at a level of only 1 µmol g^−1^ FW, fructose at a level of only 0.5 µmol g^−1^ FW but sucrose at a level of 27 µmol g^−1^ FW, which was about 5-fold higher than in WT plants (Fig. 8B). Under cold, mutant line #5 contained glucose at a level of only 10 µmol g^−1^ FW, fructose at a level of only 3 µmol g^−1^ FW and sucrose at a level of 18 µmol g^−1^ FW (Fig. 8B).

Accumulation of sugars is a prerequisite to tolerate sub-zero temperatures (Alberdi and Corcuera, 1991). Since both *amiR vi1-2* lines exhibited a markedly altered sugar composition, when compared to WT plants, we conducted electrolyte leakage assays (Patzke *et al*., 2019; Klemens *et al*., 2014; Klemens *et al*., 2013) to compare frost tolerance properties of WT and *amiR vi1-2* mutants. To this end, we transferred WT and mutant plants after four weeks of growth at 22°C for four days into the cold (4°C), to induce molecular and physiological acclimation to low temperatures. After this phase, leaves were taken, frozen at −6°C in water and, after re-thawing, ion release from disrupted cells was quantified via electro-conductivity of the supernatant. In comparison, we quantified ion release from leaves of non-acclimated plants, kept at 22°C.

Freezing of non-acclimated WT and two *amiR vi1-2* lines led to an ion loss of about 82–86% from leaves of all lines (Fig. 8C). Acclimation to low temperature correlated with a markedly improved freezing tolerance as indicated by an ion release of only about 19%, which was however independent of the tested genotype (Fig. 8C).

## Discussion

Since sucrose fulfills various roles in plant development and stress response (Rolland *et al*., 2006; Hanson and Smeekens, 2009; Teng *et al*., 2005) it appears likely that its intracellular homeostasis must be tightly regulated (Pommerrenig *et al*., 2018). This assumption gained direct support by the recent discovery of a chloroplast located sucrose exporter pSuT, which is involved in fine control of the onset of flowering and frost tolerance of Arabidopsis (Patzke *et al*., 2019).

However, mesophyll vacuoles play a prime function in the control of intracellular sucrose homeostasis, because these organelles occupy most of the volume in plant cells and represent the major storage compartment for sucrose and other sugars (Martinoia *et al*., 2007). Interestingly, although the presence of sucrose in chloroplasts, the cytosol and the vacuole has been demonstrated since decades (Heber, 1957; Gerhardt *et al*., 1987), no detailed analysis on the impact of sucrose import into vacuoles and subsequent sucrose hydrolysis in mesophyll vacuoles has been carried out so far. Thus, to influence the intracellular sucrose compartmentation in mesophyll cells we followed two routes. Firstly, we overexpressed the tonoplast located sucrose importer TST2.1 from sugar beet (Jung *et al*., 2015). Secondly, we generated Arabidopsis mutants with decreased vacuolar invertase activity. The detailed description of both types of mutants provided novel insights into the impact of sucrose import and sucrose hydrolysis in plant vacuoles for developmental processes of Arabidopsis.

### Vacuolar sucrose accumulation is not controlled by the rate of sucrose import

Previously it has been shown that *Bv*TST2.1 is a highly active, proton driven vacuolar sugar importer, able to pump sucrose against a steep concentration gradient into the vacuole from sugar beet tap-root cells (Jung *et al*., 2015). To our surprise, overexpression of *BvTST2.1* in an Arabidopsis mutant lacking the endogenous vacuolar TST type sugar loaders TST1 and TST2 led to an increase of the monosaccharides glucose and fructose, while sucrose was not affected (Fig. 1A). According to this observation we speculated that luminal hydrolysis of vacuolar located sucrose by invertase might be responsible for the inability of *BvTST2.1* overexpressor mutants to accumulate sucrose above the levels observed in WT plants and the *tst1-2* line.

To verify this assumption we exploited two alternative systems. Firstly, overexpression of the vacuolar sugar transporter *BvSUC4*, representing the closest sugar beet homolog to the vacuolar sucrose exporter SUC4 from Arabidopsis (Schneider *et al*., 2012), provoked opposite effects on the total sugar levels as seen for *BvTST2.1* overexpressors, namely lower monosaccharide levels than seen in the WT background genotype (Fig. 1B). Secondly, a transient ectopic expression of *BvTST2.1* in *Nicotiana benthamiana* leaves provoked strongly increased levels of total monosaccharides (Fig. 2B,C). However, this increase in monosaccharide levels was completely absent in leaf tissue expressing both, the invertase inhibitor protein *NbVIF* and *BvTST2.1* (Fig. 2C). Thus, all three different experimental approaches provided evidence that in leaf mesophyll cells vacuolar invertases contribute substantially to the overall sucrose hydrolysis. Moreover, we can now conclude that in Arabidopsis the activity of the vacuolar invertase, and not the rate of sucrose import, limits the sucrose storing capacity of leaves (Figs 1A, 3C). Latter conclusion is in line with comparative observations on sucrose storing organs and source leaves. This is, because in explicit sugar storing tissue like sugar beet tap roots the vacuolar invertase activity is negatively correlated with sucrose levels (Leigh *et al*., 1979).

Interestingly, Arabidopsis mutants exhibiting decreased total vacuolar invertase activity (Fig. 3A,B) showed both, substantial decrease of glucose and fructose (Fig. 3C), while sucrose levels in the vacuolar lumen were increased (Fig. 4C). These observations indicate that the endogenous vacuolar sugar importer TST from Arabidopsis, which is present in *amiR vi1-2* mutants, transports under *in vivo* conditions not only monosaccharides but also sucrose. This conclusion receives direct support by electrophysiological studies conducted on recombinant expressed *At*TST1 protein (Schulz *et al*., 2011). Since a similar substrate spectrum has also been reported for storage parenchyma cell-located sugar beet homolog *Bv*TST1 (Jung *et al*. 2015), we propose that TST proteins located in leaves are able to transport both, monosaccharides and sucrose.

### Decreased VI activity affects seed development, yield, and plant survival in the dark

Mutants, suppressing VI1-2 activity exhibited impaired shoot development in the dark (Fig. 5A,B), which concurs with previous data documenting that sugars act *per se* as positive effectors of early hypocotyl growth in the dark (Zhang *et al*., 2015). It is obvious, that etiolated seedlings from both *amiR vi1-2* lines contained markedly lower glucose levels when compared to corresponding WT seedlings, which is most likely due to a blockage of sucrose hydrolysis in mutant vacuoles (Fig. 5C). We propose for several reasons that glucose accumulation in WT seedlings has to occur in the cytosol to be functional as a positive trigger for etiolated growth. Firstly, sugar stimulation of early plant growth depend upon the presence of functional brassinosteroid signaling (Peng *et al*., 2018), which is located in the cytosol (Planas-Riverola *et al*., 2019). Secondly, the expression of the gene coding for the vacuolar glucose exporter ERDL6 is increased under conditions of low sugar availability, to promote sugar release into the cytosol (Poschet *et al*., 2011). Thirdly, the cytosolic sugar sensing HEXOKINASE1 is critical for glucose-induced etiolated growth of Arabidopsis (Zhang and He, 2015). Moreover, that monosaccharides released from sucrose hydrolysis must be present in the cytosol is also indicated by analyses of dark growth of Arabidopsis seedlings lacking the cytosolic invertase activity. Latter mutants exhibit, similar to *amiR vi1-2* plants (Fig. 5A,B) impaired shoot development (Barnes and Anderson, 2018), pointing to a synergistic function of both invertase isoforms (cytosolic and vacuolar) for carbon supply under conditions of low energy. Thus, our data raised on *amiR vi1-2* seedlings explain how glucose is released into the cytosol of etiolated plants. Obviously, sufficient vacuolar invertase activity is required for Arabidopsis development under dark conditions, in which an efficient conversion of stored lipids into energy providing sugars is mandatory.

Temporal darkness of single leaves of larger parts of a plant are naturally occurring situations and are responded by a plant with a complex, phytochrome-dependent shade (dark) avoidance program (Martínez-García *et al*., 2014), allowing to direct plant growth towards light. However, for such growth direction, energy is required and rapid mobilization of various storage compounds, *inter alia* sugars, is one of the prime metabolic rearrangements (Law *et al*., 2018). Thus, the observation of an increase of VI activity upon onset of an extended dark period concurs with these dark induced metabolic modifications (Fig. S3A,B) indicates that sucrose hydrolysis in the vacuolar lumen is critical for dark response and survival. In this respect it was surprising to see that both *amiR vi1-2* lines exhibited an unexpected tolerance against an extended exposure to darkness (Fig. 7A,B). Such optimized tolerance correlates with increased sucrose levels in both *amiR vi1-2* mutants when compared to WT plants, especially after three days in complete darkness, leading to higher monosaccharide availability when compared to WT plants (Fig. 7C,D). Accordingly, minimal levels of cellular sugars seem to prevent the completion of the dark induced senescence program. This conclusion gains support by similar observations on leaves from various plant species fed with sugars, and incubated in extended darkness (Thimann *et al.*, 1977; Chung *et al*., 1997).

Thus, the question arises why in WT plants the transcript level of *VI2* is not downregulated in total darkness, although such molecular response might extend the time of survival under latter conditions. However, normally, as a photoautotroph organism, plants are not adapted to survive very long dark phases. Instead, they must mobilize energy reserves rapidly to avoid shade by growth to ensure survival and reproduction (Morelli and Ruberti, 2002). Obviously, the ecologic reaction of plants is to allow rapid growth for entering light, and not to survive darkness for extremely long phases. Interestingly, potato mutants with impaired activity of a plasma membrane located sugar transporter *St*SUT4 also exhibit decreased shade-avoidance (Chincinska *et al*., 2008). In summary, a controlled carbohydrate homeostasis, involving sufficient vacuolar sucrose cleavage, is an element for proper reaction upon dark stress (Chincinska *et al*., 2008).

### Impaired seed yield in mutants with reduced VI activity

The development of sink tissues depends on an efficient conversion of sucrose, or in some species imported polyols, into suitable storage products like starch, triacylglycerols (lipids), or storage proteins (Baghalian *et al*., 2014). Given that vegetative parts of both *amiR vi1-2* lines are highly similar to WT plants in size (Fig. 6A,B), it was surprising to see that corresponding seeds from mutant plants exhibit decreased seed weight, correlating with lower lipid levels (Fig. 6C,D). Since the size and mass of vegetative rosettes, responsible for source activity, of both mutant lines are similar to WT plants (Fig. 6A), we attribute decreased seed size and lipid content of mutants to specific function of the vacuolar invertase in carbon filling into siliques and developing seeds. This assumption gains support by research on other species, showing that decreased vacuolar invertase activity in muskmelon or rice promote decreased fruit and seed size, respectively (Yu *et al*., 2008; Lee *et al*., 2019). Moreover, as seen in siliques from *amiR vi1-2* mutants (Fig. 6E), decreased vacuolar invertase in developing rice seeds led to impressively increased sucrose/monosaccharide ratio (Lee *et al*., 2019). It has been shown that developing Arabidopsis seeds can use both, sucrose and monosaccharides for their development, but especially in the phase of early seed development the sum of glucose and fructose exceeds the sucrose levels significantly (Hill *et al*., 2003). Thus, it appears likely that founding of further storage cells is stimulated by vacuolar invertase activity, since glucose, but not sucrose, serves as a cellular signal promoting cell proliferation (Wang and Ruan, 2013).

### Altered vacuolar sucrose homeostasis does not affect frost resistance

The observation that cold exposure leads to a marked down-regulation of mRNAs coding for VI1 and VI2 (Fig. 8A) is in line with previous findings that various Arabidopsis accessions tune down vacuolar invertase activity after exposure to low temperature (Nägele and Heyer, 2013). Thus, that cold-treated Arabidopsis plants accumulate not only sucrose but also glucose and fructose in the vacuole (Nägele and Heyer, 2013) is most likely due to both, decreased sucrolytic activity in the vacuolar lumen and increased activity of vacuolar monosaccharide import, accompanied by decreased glucose and fructose export (Wormit *et al*., 2006; Klemens *et al*., 2014; Poschet *et al*., 2011). In addition, it has been proposed that cold-induced shift of total sucrose cleavage from the cytosol into the vacuole might be required for a cytosolic accumulation of sucrose, known to act as an agent against frost injuries (Weiszmann *et al*., 2018a).

Interestingly, both *amiR vi1-2* lines exhibit markedly increased sucrose levels in leaves and decreased concentrations of monosaccharides when compared to WT plants (Fig. 8B). Given, that frost tolerance of both *amiR vi1-2* mutants is similar to WT plants (Fig. 8C), we can thus state that increased vacuolar sucrose levels fully compensate for low monosaccharide concentrations. It will be interesting to check whether *amiR vi1-2* lines exhibit, when compared to WT plants, other alterations in response to cold temperature. For example, plants unable to synthesize raffinose do not show decreased frost tolerance, while protection of photosynthetic capacity during frost is impaired in these mutants (Knaupp *et al*., 2011; Zuther *et al*., 2004). Obviously, decreased vacuolar invertase activity does not impair cold hardiness. Thus, further investigations have to be carried out to understand the evolutionary advantage of monosaccharide accumulation in cold acclimated Arabidopsis plants.

## Abbreviations

amiRNA: artificial microRNA
At: Arabidopsis thaliana
Bv: Beta vulgaris
Nb: Nicotiana benthamiana
TST: tonoplast sugar transporter
VI: vacuolar invertase
WT: wild-type

## Acknowledgements

We thank the LMUexcellence Junior Researcher Fund (Nägele lab). This work was financially supported by the Deutsche Forschungsgemeinschaft (DFG) in the frames of the IRTG1831 and TRR175 (Nägele lab and Neuhaus lab).

## Supplementary data

Table S1. Primer sequences for cloning.

Table S2. Primer sequences for expression analyses.

Fig. S1. Generation of *BvTST2.1* overexpressors in the genetic background of a *tst1-2* double mutant under control of a constitutive promoter.

Fig. S2. Generation of *BvSUC4* overexpressors in the wild-type background under control of a constitutive promoter.

Fig. S3. Molecular and biochemical characterization of dark treated wild-type plants.

