## Supplementary Tables 1-2 for "Vacuolar sucrose homeostasis is critical for development, seed properties and survival of dark phases of Arabidopsis"

**Supplementary Information**

**Tables S1-S2**

| **Table S1.** Primer sequences for cloning. | | |
| --- | --- | --- |
| Purpose | Primer | Sequence (5’ to 3’) |
| Cloning of amiRNA fragment | AtVI1+2_Ia_fw | gaTAAGGATGAATAAAAGCACGGtctctcttttgtattcc |
|  | AtVI1+2_IIa_rv | gaCCGTGCTTTTATTCATCCTTAtcaaagagaatcaatga |
|  | AtVI1+2_IIIa_fw | gaCCATGCTTTTATTGATCCTTTtcacaggtcgtgatatg |
|  | AtVI1+2_IVa_rv | gaAAAGGATCAATAAAAGCATGGtctacatatatattcct |
|  | A_miR_fw | CTGCAAGGCGATTAAGTTGGGTAAC |
|  | B_miR_rv | GCGGATAACAATTT ACACAGGAAACAG |
| Cloning into Gateway™ system | pR300_GW_fw | GGGGACAAGTTTGTACAAAAAAGCAGGCTCCTCGAGGTC  GACGGTATC |
|  | pR300_GW_rev | GGGGACCACTTTGTACAAGAAAGCTGGGTGGCCGCTCTA  GAACTAGTGGA |

| **Table S2.** Primer sequences for expression analyses. | | |
| --- | --- | --- |
| Gene | Primer | Sequence (5’ to 3’) |
| *BvTST2.1* | qPCRfwTST2.1 | AAAGATGAACACCACTGTGTATG |
|  | qPCRrevTST2.1 | GTCATCAGTGGCTTGCTTGTCTTG |
| *BvSUC4* | BvSUC4-f | TGACACTGACTGGATGGGTC |
|  | BvSUC4-r | CCCAGAACCCAACTTCTTGC |
| *AtVI1* | AtVI1_qPCR_fw | GGTTGGTCTTCTGTTCAGGGCATC |
|  | AtVI1_qPCR_rev | ACCACAGTCCCTGGTCCAATAGTC |
| *AtVI2* | AtVI2_qPCR_fw | ACGACCAGAACAAGGGTCGAAG |
|  | AtVI2_qPCR_rev | AACGGTTCTTGGGATACCCTGGAG |
| *AtPP2A* | AtPP2A_qPCR_fw | TAACGTGGCCAAAATGATGC |
|  | AtPP2A_qPCR_rv | GTTCTCCACAACCGCTTGGT |
| *AtSAND* | AtSAND_qPCR_fw | aactctatgcagcatttgatccact |
|  | AtSAND_qPCR_rv | tgattgcatatctttatcgccatc |
