## Supplementary Figures 1-3 for "Vacuolar sucrose homeostasis is critical for development, seed properties and survival of dark phases of Arabidopsis"

**Supplementary Information**

**Figures S1-S3**

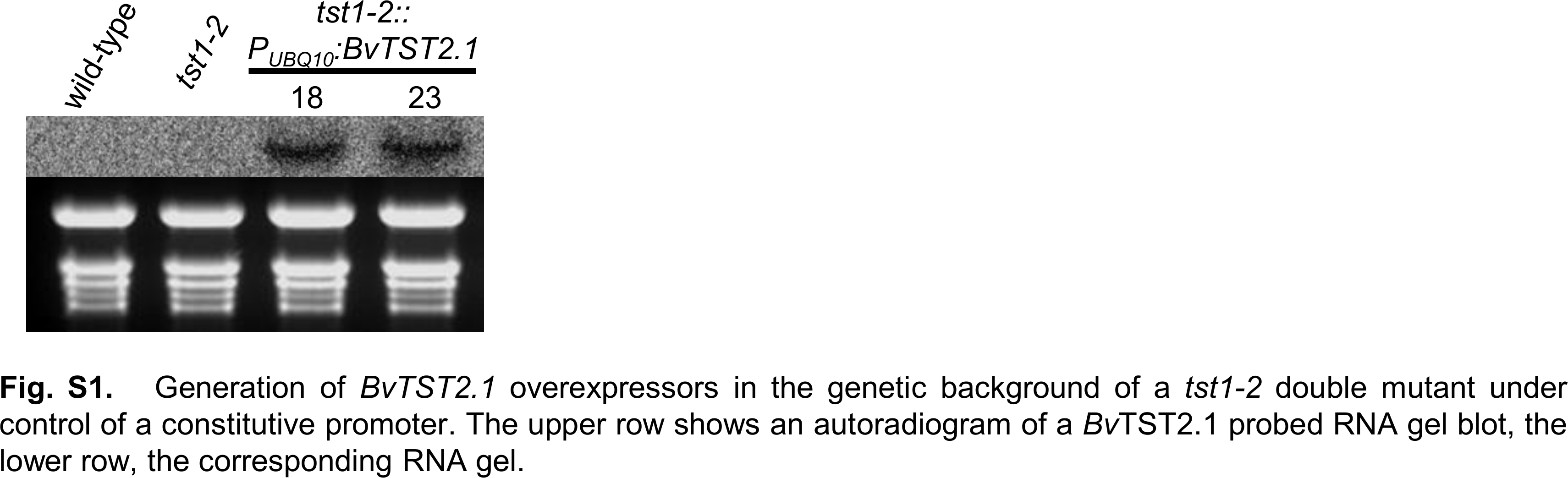

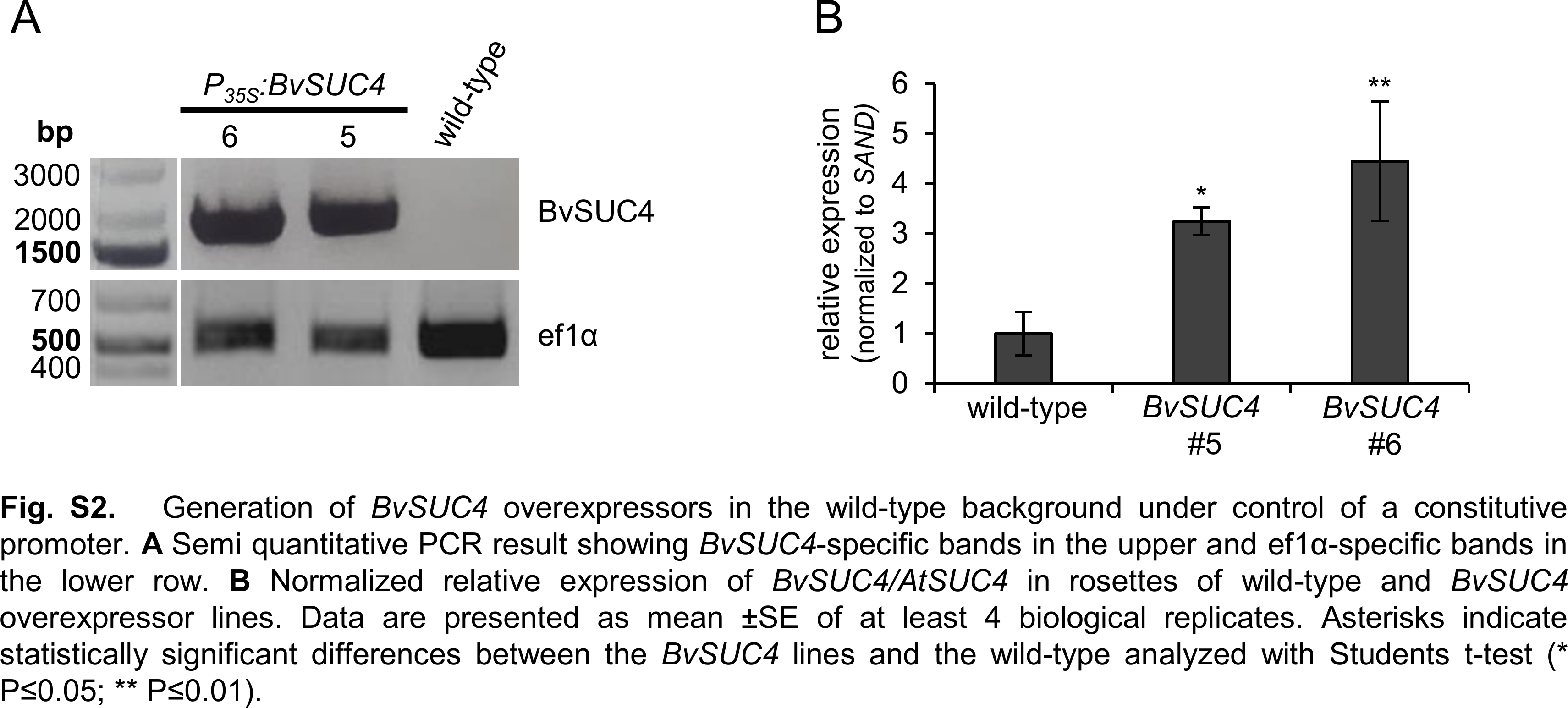

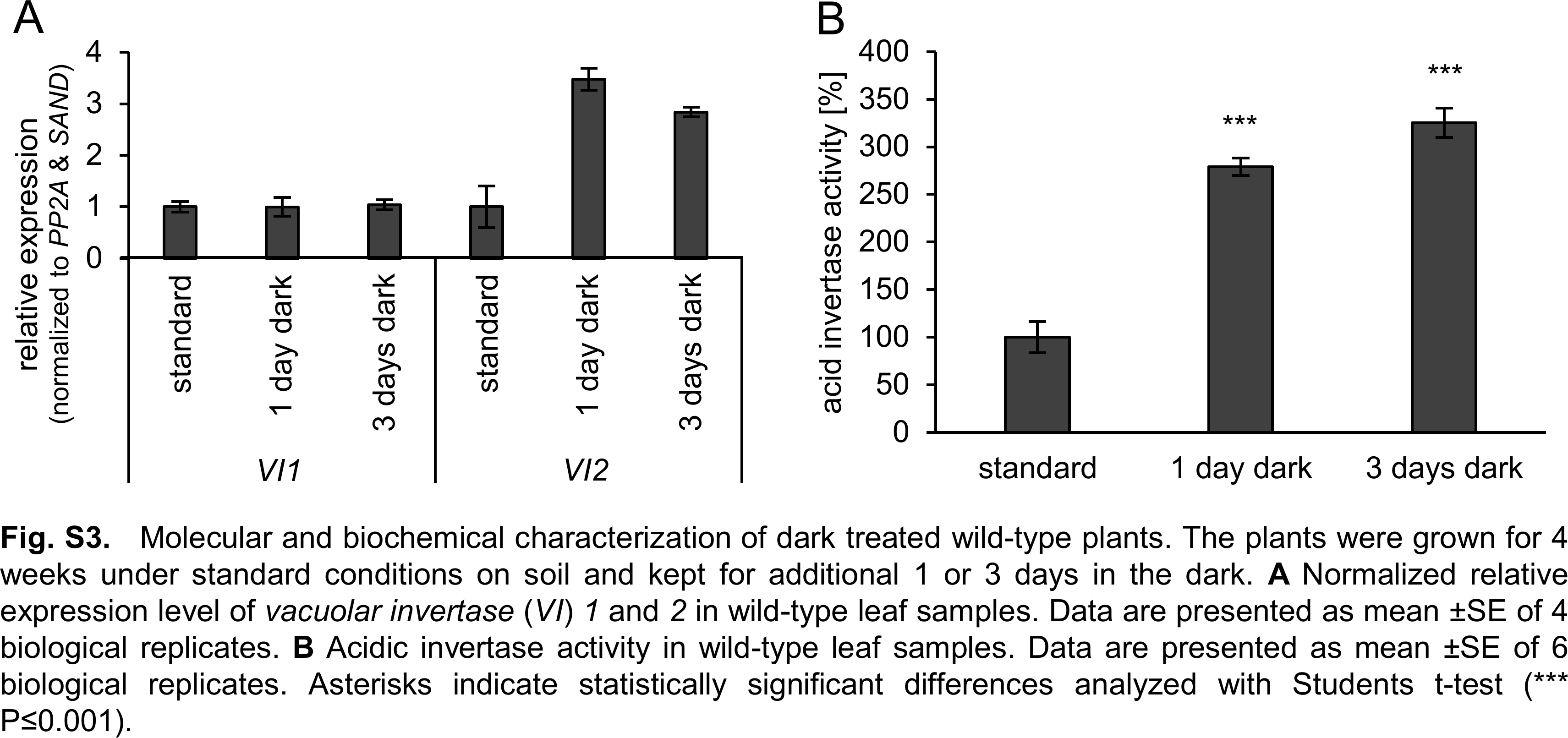
